# What is a Differentially Expressed Gene?

**DOI:** 10.1101/2025.01.31.635902

**Authors:** Franziska Hoerbst, Gurpinder Singh Sidhu, Melissa Tomkins, Richard J. Morris

## Abstract

The concept of ‘Differentially Expressed Genes’ (DEGs) is central to RNA-Seq studies, yet their identification suffers from reproducibility issues. This is largely a consequence of the inherent biological and technical variation that cannot be captured with small numbers of replicates. When thresholds for p-values and log_2_ fold changes are introduced, this variability can propagate an incomplete description of the data, leading to differing interpretations. Here, we compare traditional binary DEG classification with a rank-based method, grounded in Bayesian statistics, using a published yeast dataset comprising over 40 replicates. This analysis reveals how the choice of thresholds and number of replicates results in discrepancies between studies and potentially interesting genes being overlooked. Furthermore, by comparing wild-type with wild-type samples, we show how variability in gene expression can be mistaken for differential expression. Evaluating current practices for navigating the accuracy-error trade-off in the search for differentially expressed genes leads us to advocate rank-based methods and Bayesian statistics to mitigate the limitations of binary classifications and communicate uncertainty.

## 1 Introduction

RNA sequencing (RNA-Seq) has revolutionized molecular biology, providing a snapshot of RNA abundances in cells or tissues and thus offering unprecedented opportunities to study transcriptomic changes in response to environmental stimuli or treatments. The technology and the associated data processing tools have advanced significantly over the years and continue to be developed and refined, improving the reliability and quantification. Nevertheless, reproducibility issues have been reported, particularly in the identification of ‘Differentially Expressed Genes’ (DEGs) [1]–[4]. To explore the source of these problems, we recently re-examined the statistical foundations of differential gene expression analysis [5].

Although the question of how experimental variation can impact the accuracy of RNA-Seq analysis is not new, previous studies largely focussed on finding the best analysis tool and/or adequate number of biological replicates to optimise accuracy [1], [6]–[16]. In this manuscript, we build on these studies to revisit the underlying statistical framework of differential gene expression analysis and identify the factors that contribute to conflicting results. Traditional methods often introduce thresholds, which can, combined with technical and biological variation, lead to inconsistent results between studies [6]–[16]. DEGs are typically determined based on two criteria: (1) p-values below a set threshold and (2) absolute log_2_ fold change above a certain threshold (both for normalised read counts between the two conditions) which can be calculated using many of the available software packages [1]–[4], [6]–[16].

Here, we explore how variability in gene expression, the number of biological replicates, and the choice of statistical tools affect the identification of DEGs. Our analysis is based on a series of experiments using published yeast RNA-Seq data [1]. We compare two popular RNA-Seq analysis packages – *DESeq2* and *edgeR* – with a new Bayesian framework, *bayexpress*. We previously showed that Bayes factors can be used to rank genes based on statistical evidence for expression change, *BF*_21_, and for evaluating the consistency across replicates, *BF*_*k*1_. The two in combination can be used to curate lists of DEG candidates for further analysis [5].By showing where different approaches agree and where they disagree, we demonstrate how the choice of thresholds for binary classification (DEG? yes/no) can impact the accuracy of the analysis. The findings challenge the widely used classification criteria and highlight the need for more sophisticated and consistent approaches to DEG analysis that account for data availability, data variability, and reduce the reliance on thresholds.

## 2 Results

### 2.1 Fold change cut-offs increase the number of false negatives

Fold change cut-offs are frequently applied with the goal of decreasing the number of potential false positives and identifying the ‘most significant’ gene expression changes. To reduce the list of genes of interest, this remains a valid approach. However, introducing thresholds, comes at the cost of missing potentially interesting genes that have undergone relevant changes below the chosen cut-off. For example, Figure 1 shows a representative case of a gene that would be missed. Naturally, without knowledge of which expression changes for each gene have downstream consequences, we are not able to identify actual true or false positives, i.e. we lack the ground truth. We can, however, point at those genes that mirror the concept of potential ‘false negatives’ to demonstrate the impact of choosing thresholds in current analyses. Therefore, we term some example genes (Figure 1) as putative ‘false negatives’ as they demonstrate potentially interesting behaviour, but would not qualify as DEG and, hence, be excluded from further analyses. Since the biological impact of an expression change is not necessarily correlated with its magnitude, a gene with a small change in expression could potentially have larger consequences than a gene that doubles its expression (log_2_ fold change > 1). Some measure of confidence in the assignment is, therefore, helpful to express uncertainty and guide decisions.

**Figure 1:**
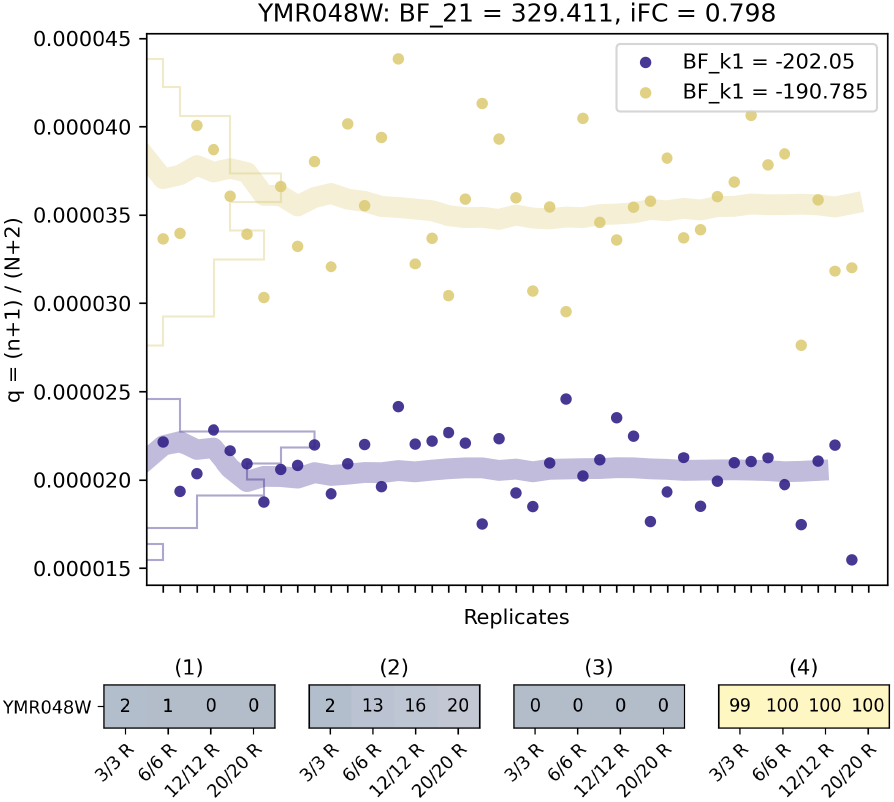
Genes with small fold-changes in expression can be identified using Bayes factors. The example, YMR048W, shows a gene for which the Bayes factor for differential expression (*BF*_21_) [5] reports strong evidence for change between the WT yeast (purple) and SNF2-mutant (yellow). DEG definitions using a log fold change cut-off of above 1 do not reliably classify this gene as a DEG. YMR048W, also known as CSM3, is required for stable replication fork pausing. Shown are the inferred expression rates, *q*, across 42 WT and 44 SNF2-mutant bulk, yeast culture RNA-Seq replicates [1]. The fine lines are density histograms, obtained by projecting the points on the y-axis, and the thick lines are estimated means of *q*, updated with each replicate. We calculated a log Bayes factor (*BF*_21_) and inferred log_2_ fold change (*iFC*), shown at the top, using all 42/44 replicates. The bottom plots show the number of times (out of 100) the gene was identified as a DEG, for randomly sampled datasets with 3, 6, 12 and 20 replicates. We examine four DEG criteria: (1) *BF*_21_ > 1, and inferred | *iFC* | > 1; (2) *edgeR*: p-value *<* 0.05 and | log_2_ fold change | > 1; (3) *DESeq2* : p-value *<* 0.05 and | log_2_ fold change | > 1; (4) *BF*_21_ > 1, no log_2_ fold change cut-off. The (log) Bayes factors for expression consistency, *BF*_*k*1_, are given as insets in boxes [5]. A negative log *BF*_*k*1_ supports all replicates being consistent with one expression rate.

We identified 5437 genes (76% of the genome) that show notable gene expression differences (*BF*_21_ > 1) in the 42 replicate WT and 44 replicate SNF2-mutant bulk, yeast culture RNA-Seq data. SNF2 is a gene involved in chromatin remodelling and transcription regulation of polymerase II promoters, explaining the sweeping changes we observe. In Table 1 and Figures 10, 11, and 12 in the Appendix we present validated SNF2 targets and discuss for which we can or cannot find evidence in the RNA-Seq data. We include several other example genes in a ‘Differential Gene Expression Plot Collection’ in the Appendix. Other genes can be explored using the resources on our GitHub.

**Table 1:** Evidence for expression changes of SNF2 targets in the RNA-Seq data. The deletion of SNF2 in the yeast genome introduces far-reaching changes in the transcriptome [27]–[30]. We evaluated the evidences for gene expression changes of a hand-curated list of direct SNF2-targets in the RNA-Seq data. We show Bayes factors (*BF*_2_1) alongside inferred log_2_ fold change and further information from the yeast genome data bank. We plotted gene expression probabilities between wild-type and mutant for those target genes in Figures 10, 11, and 12. Note, that ‘YHR053C’ and ‘YHR055C’ do not have any reads mapping to them.

| Target | Target Systematic Name | BF<sub>21</sub> | IFC | Happens During | Regulator Type | Direction | Happens During | Evidence | Reference |
| --- | --- | --- | --- | --- | --- | --- | --- | --- | --- |
| CUP1-1 | YHR053C | -8.274 | -0.138 | cellular response to copper ion | chromatin modifier | positive | cellular response to copper ion | quantitative reverse transcription polymerase chain reaction evidence | Wimalaratna RN, et al. (2014) PMID:24471920 |
| CUP1-2 | YHR055C | -8.274 | -0.138 | cellular response to copper ion | chromatin modifier | positive | cellular response to copper ion | quantitative reverse transcription polymerase chain reaction evidence | Wimalaratna RN, et al. (2014) PMID:24471920 |
| ECM4 | YKR076W | 3095.500 | 1.495 | cellular response to cadmium ion | chromatin modifier | negative | cellular response to cadmium ion | reverse transcription polymerase chain reaction evidence | Sahu RK, et al. (2020) PMID:31846549 |
| GTO1 | YGR154C | 10.097 | -0.207 | cellular response to cadmium ion | chromatin modifier | negative | cellular response to cadmium ion | reverse transcription polymerase chain reaction evidence | Sahu RK, et al. (2020) PMID:31846549 |
| GTT2 | YLL060C | 2390.104 | 1.552 | cellular response to cadmium ion | chromatin modifier | negative | cellular response to cadmium ion | reverse transcription polymerase chain reaction evidence | Sahu RK, et al. (2020) PMID:31846549 |
| HO | YDL227C | 537.817 | -1.742 |  | chromatin modifier | positive |  | beta-galactosidase reporter gene assay evidence | Richmond E and Peterson CL (1996) PMID:8871545 |
| HSP26 | YBR072W | 56.779 | 0.401 | cellular response to cadmium ion | chromatin modifier | negative | cellular response to cadmium ion | reverse transcription polymerase chain reaction evidence | Sahu RK, et al. (2020) PMID:31846549 |
| INM1 | YHR046C | 8.300 | -0.136 | cellular response to inositol | chromatin modifier |  | cellular response to inositol | chromatin immunoprecipitation-qPCR evidence | Zhang L and Di J (2014) PMID:25211324 |
| SPL2 | YHR136C | 8284.989 | -4.401 |  | chromatin modifier | positive |  | northern blot evidence | Ellison MA, et al. (2019) PMID:31092540 |
| SUC2 | YIL162W | 110.225 | -0.195 |  | chromatin modifier | positive |  | enzymatic activity assay evidence | Richmond E and Peterson CL (1996) PMID:8871545 |
| TPK1 | YJL164C | 231.277 | 0.357 | cellular response to heat | chromatin modifier |  | cellular response to heat | quantitative reverse transcription polymerase chain reaction evidence | Reca S, et al. (2020) PMID:32599085 |
| VTC3 | YPL019C | 17426.468 | -2.211 |  | chromatin modifier | positive |  | northern blot evidence | Ellison MA, et al. (2019) PMID:31092540 |

### 2.2 Genes with inherently variable expression can lead to false positives

We conducted a computational experiment to investigate the impact of data variability on the identification of DEGs. We created 100 datasets for 3 and 10 replicates by sampling from a pool of 42 wild-type replicates. For each gene, we calculated Bayes factors for differential expression (*BF*_21_), which identified hundreds or thousands of DEGs, depending on chosen cut-offs, Figure 2. Such ‘significant DEGs’ found in this control experiment can be viewed as ‘false positives’. Our analysis revealed that while most DEGs exhibited moderate log_2_ fold changes, a surprising number of genes, particularly in datasets with only 3 replicates, displayed noteworthy deviations. We observe high Bayes factors, which demonstrate that strong evidence for changes in expression can be found even within a control experiment. The findings presented here serve as a compelling reminder of the importance of accounting for data variability, and exercising prudence in the interpretation and communication of differential gene expression results, particularly in scenarios with limited numbers of replicates.

**Figure 2:**
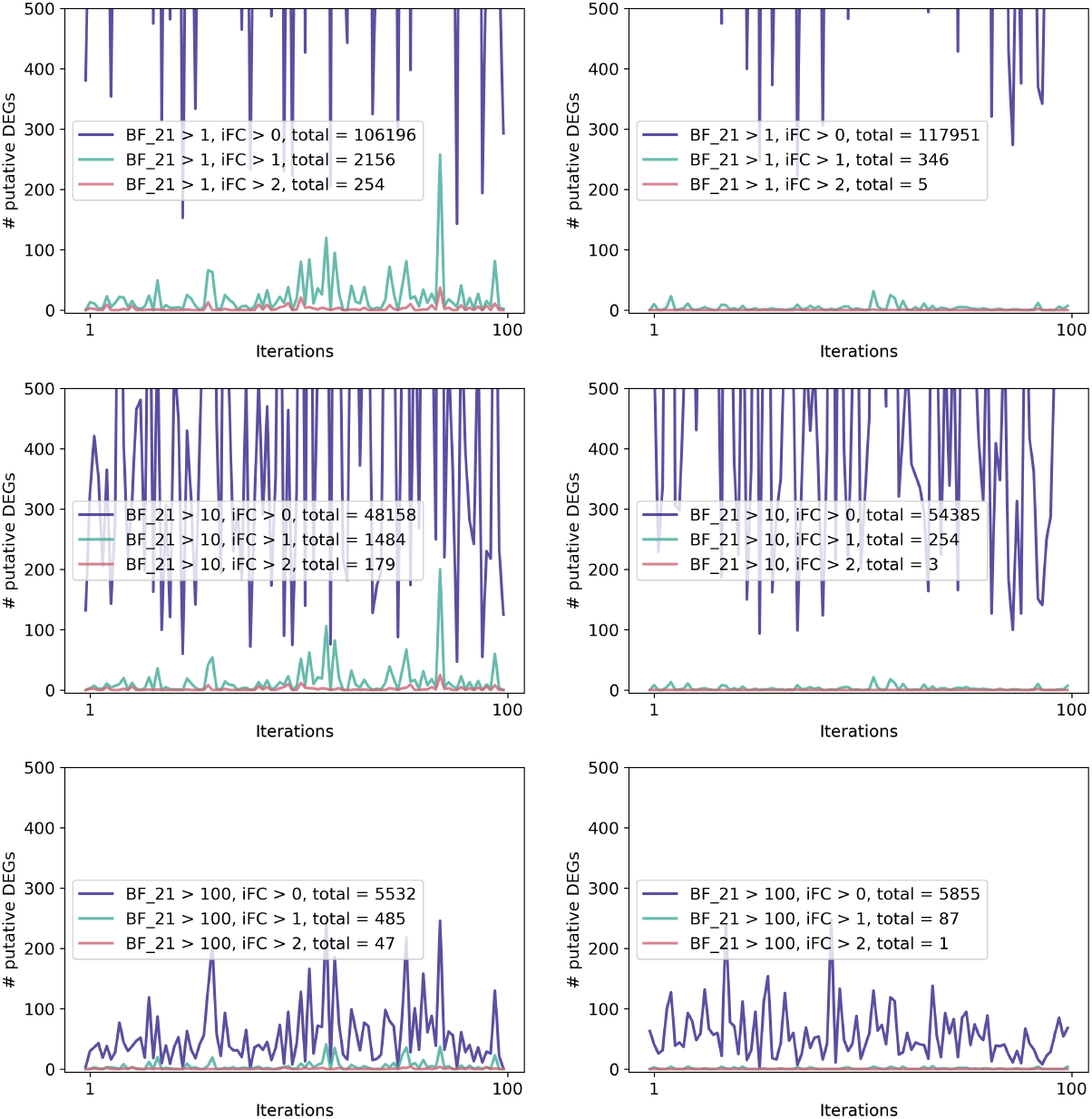
A control experiment comparing wild-type with wild-type data demonstrates how variability can lead to the identification of expression changes that may incorrectly be associated with a perturbation (DEG). These plots show the number of putative DEGs (y-axis) identified in 100 randomly sampled datasets (x-axis) from 44 replicates in a yeast experiment. We consider 7126 genes and examine three different criteria denoted by purple (no log_2_ fold change cut-off), green (| log_2_ fold change | > 1) and red (| log_2_ fold change | > 2) lines. The first row is using a Bayes factor (*BF*_21_) cut-off of 1, the second row uses a Bayes factor (*BF*_21_) cut-off of 10 and the third row uses a Bayes factor (*BF*_21_) cut-off of 100. In the left column the number of replicates in each sampled data set is 3, in the right column it is 10. Interestingly, the purple criteria (no log_2_ fold change cut-off) has a higher total number of putative DEGs in the 10 replicate experiment compared to 3 replicates. The third row shows DEGs with Bayes factor > 100, which can be interpreted as extremely strong evidence.

### 2.3 Highly variable genes can masquerade as DEGs

Finding DEGs in the control experiment with replicates from only one condition (wild-type) led us to ask how many genes have sufficient variability in read counts to be identified as DEGs. Expression variability has been described in detail [2]– [4], [7]–[10], [17]–[20], yet how much of the variability is of technical or biological nature remains challenging to discern [21]–[24].

The large number of replicates in the data set allowed us to identify genes with consistently high variability. We calculated Bayes factors, *BF*_*k*1_, as a function of the number of replicates (2 to 42 for WT, and 2 to 44 for mutant). We repeat this process 100 times for both WT and mutant, shuffling the set of replicates at every iteration, Figure 3. The number of identified genes converges with increasing numbers of replicates. For a gene to be considered what we term ‘consistently inconsistent’ after analysing *k* replicates, we require *BF*_*k*1_ > 1. Out of 7126 genes in yeast, the number of genes that were identified as consistently inconsistent in at least one iteration were 1633 and 922 in wild-type and mutant, respectively. This is equivalent to 23% and 13% of the genome. We found 1426 and 765 genes to be consistently inconsistent in all 100 iterations in wild-type and mutant, respectively. In Figure 9 in the Appendix we show example genes of this set, their expression across replicates, and the DEG classification by *edgeR, DESeq2* and *bayexpress*.

**Figure 3:**
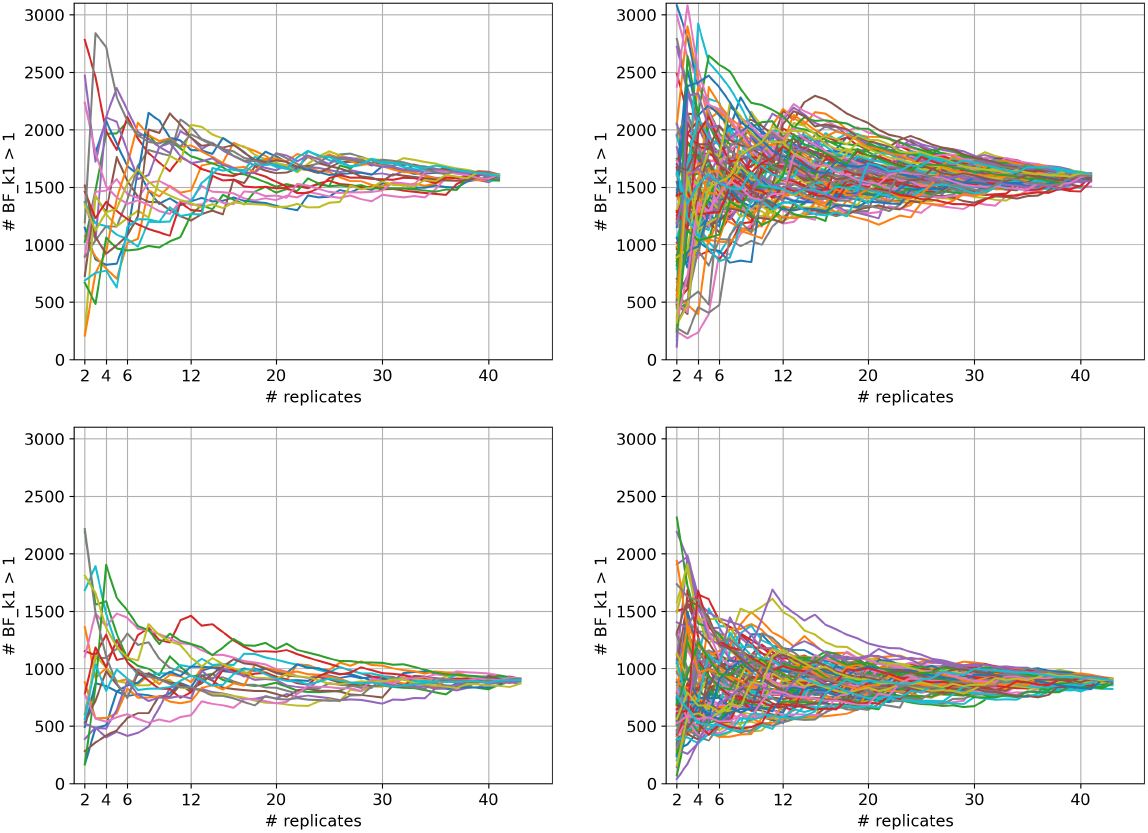
Genes with variable expression can be mistaken for DEGs. High numbers of replicates allow for the variability in expression to be robustly assessed. Using a yeast dataset with over 40 replicates, we identified approximately 1600 consistently inconsistent genes in the WT (top) and around 900 in the SNF2-mutant (bottom). These genes are prone to misclassification as DEG in studies with insufficient replicates (Supplementary Figure 13). We calculated Bayes factors (*BF*_*k*1_) for the consistency of gene expression in RNA-Seq with increasing numbers of replicates. We shuffled the replicates and repeated the process 100 times (right), each repetition being represented in a different colour. For visualisation purposes we include a plot with only 20 repetitions on the left. The y-axis denotes the number of genes that have been identified as inconsistent across replicates. The criterion is a Bayes factor for consistency *BF*_*k*1_ > 1. The x-axis depicts the number of replicates we take into account. Interestingly, we notice a different number of inconsistent genes in WT and mutant.

## 3 Discussion

Several reports show that differential gene expression studies suffer from reproducibility issues [6]–[16]. In this study, we examine the use of different analysis frameworks, the application of fold change cut-offs, and the number of replicates considered in DEG experiments. This leads us to question current definitions of differentially expressed genes.

We recently reported that with sufficient numbers of replicates, discrepancies between results can be explained by differing DEG definitions [5]. Here, we show that common numbers of replicates, typically justified by cost-benefit considerations, are often insufficient to capture biological and technical variation. To investigate the impact of the arising incomplete description of the data, we conducted three experiments using published yeast data.

In the first experiment, we compared different statistical methods for differential gene expression analysis and learned how insufficient numbers of replicates cannot provide the necessary information to draw conclusions (especially for smaller fold changes), and may produce false positives. Introducing fold change cut-offs, on the other hand, leads to potential false negatives. This is an example of how the accuracy-error trade-off cannot be circumvented nor improved by better statistics alone, requiring other interventions. Some showcase examples are depicted and discussed in the Appendix.

In the second experiment, a control experiment comparing wild-type with wild-type data, we showed how high variability of genes in RNA-seq data across replicates can result in DEGs being identified.

In the third experiment, we investigated the prevalence of expression variability. We found 23% of genes in the WT and 13% in the SNF2-mutant to be consistently inconsistent. These genes are more likely to appear as false positives (i.e. they can explain up to 70% identified DEGs, Figure 13), especially if insufficient replicates are employed. Considering the high number of problematic genes, this raises questions about the value and insights gained by large-scale techniques if not handled carefully.

The above findings build on the assumption that investigating changes in gene expression is a binary classification problem. In contrast, we recently proposed that ranking genes using Bayes factors [5] offers an alternative to conventional binary classifications. This reduces the loss of information in the analysis and the loss of potentially interesting genes. Bayesian frameworks allow consideration of the variability of genes between replicates, and enable the communication for uncertainty.

Increasing the number of biological replicates in RNA-Seq experiments is a straight-forward yet impactful way to enhance data resolution and reliability. Previous studies have suggested that 6 to 12 replicates are necessary to achieve robust results, depending on the desired sensitivity of the analysis [1]. Maybe we need to rethink common cost-benefit considerations for choosing the number of replicates.

Finally, it is important to document and publish all results, regardless of their perceived significance. Published data and repositories provide tremendous potential for learning about genes of interest and their expression patterns and variability. Contributing to and using such resources and practices will grant a more comprehensive understanding of RNA-Seq data and facilitate the critical review and, hence, improvement of their analysis and interpretation.

## 4 Materials and Methods

The experimental and simulated data we used can be found with scripts to repeat the study and reproduce all figures on our GitHub. The introduction of Bayes factors for differential gene expression (*BF*_21_) and Bayes factors to evaluate the consistency across replicates (*BF*_*k*1_) was recently published [5].

### 4.1 Yeast data

Our package comparison was inspired by a yeast study conducted by Schurch et al. [1], [25]. They performed RNA-Seq on 42 wild-type (WT) and 44 SNF2-mutant replicates and carried out a bootstrapping study to find out how many biological replicates RNA-Seq studies need and which packages to use for the analysis. Thanks to their detailed documentation and publicly available data, we could build up on their work. We used their fully processed data downloaded from their GitHub repository. As documented in [1], [25], we worked with 42 WT and 44 mutant replicates within the original data set of 48 replicates. *Saccharomyces cerevisiae* was chosen for its well-studied and relatively small transcriptome with very limited alternative splicing. SNF2 is known to be the catalytic subunit of a chromatin remodelling complex called SWI/SNF. It forms part of a transcriptional activator and its deletion causes wide-ranging changes across the transcriptome. The evidence for gene expression changes of a hand-curated list of SNF2-targets, can be found in Table 1 and Figures 10, 11, and 12 in the Appendix. Interestingly, although the mutant is a deletion of SNF2, we find around 10 reads for this gene in all replicates.

### 4.2 Random sampling of datasets

We created 100 datasets by randomly sampling, with replacement, either 3, 6, 12, or 20 replicates from the pool of 42/44 yeast replicates. Each replicate can only appear once within a data set. We set up a pipeline to analyse the data using *edgeR, DESeq2* and our Bayesian framework *bayexpress* to compare different packages (Figures 1, 4, 5, 6, 7, 8 and 9). This data and analysis pipeline was also used for the WT-WT control experiment (Figure 2) and the identification of consistently inconsistent genes (Figure 3). All datasets and results have been stored and published, as well as all code to reproduce our findings.

**Figure 4:**
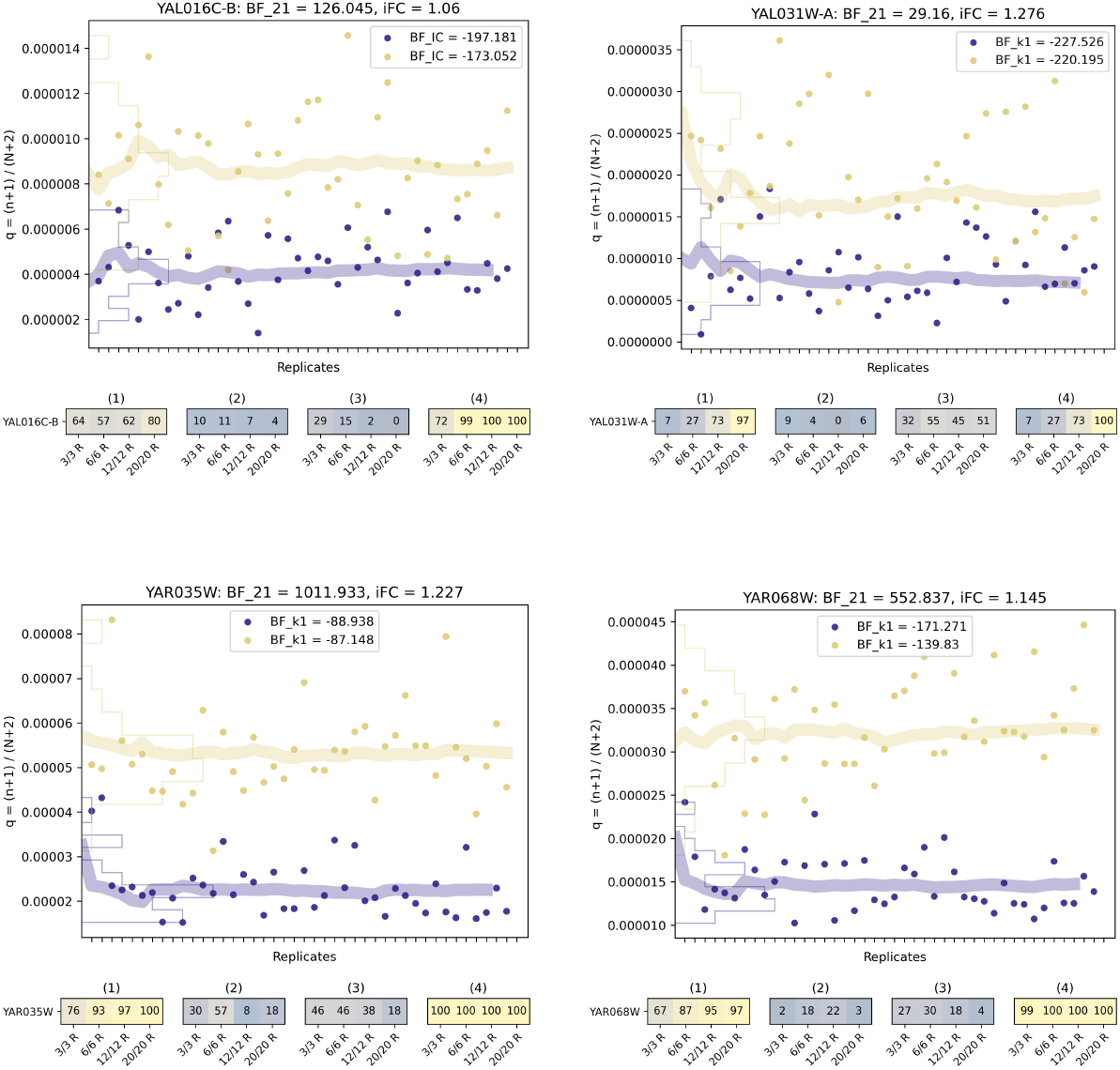
A set of example genes in yeast for which the DEG classification is sensitive to log_2_ fold change cut-offs. According to the calculated Bayes factors (*BF*_21_), we find evidence for gene expression change in the data for the four example genes. The magnitude of *BF*_21_ depends on the number of replicates. The examples gene are not reliably identified by *edgeR* and *DESeq2*. The plots at the top show inferred expression probability, *q* = (*n* + 1)*/*(*N* + 2) with *n* reads mapping to the gene of interest and *N* total reads in the experiment. We show one gene per plot across 42 WT and 44 SNF2-mutant bulk, yeast culture RNA-Seq replicates [1]. The WT is shown in purple, and the SNF2-mutant in yellow. Note the different scales of the y-axes. The fine lines are density histograms obtained by projecting the points on the y-axis, and the thick lines are estimated means of *q*, updated with each replicate. For each gene Bayes factor (*BF*_21_) and inferred log_2_ fold change (*iFC*) over all replicates are given at the top. The bottom plot shows the number of times (out of 100) the gene has been identified as a DEG, for randomly sampled datasets with 3, 6, 12 and 20 replicates. We examine four DEG criteria: (1) *BF*_21_ > 1, and inferred | *iFC* | > 1; (2) *edgeR*: p-value *<* 0.05 and | log_2_ fold change | > 1; (3) *DESeq2* : p-value *<* 0.05 and | log_2_ fold change | > 1; (4) *BF*_21_ > 1, no log_2_ fold change cut-off. The (log) Bayes factors for expression consistency, *BF*_*k*1_, are given as insets in boxes [5]. If the gene is marked* it has been identified as ‘consistently inconsistent’, see Results section and Figure 3.

**Figure 5:**
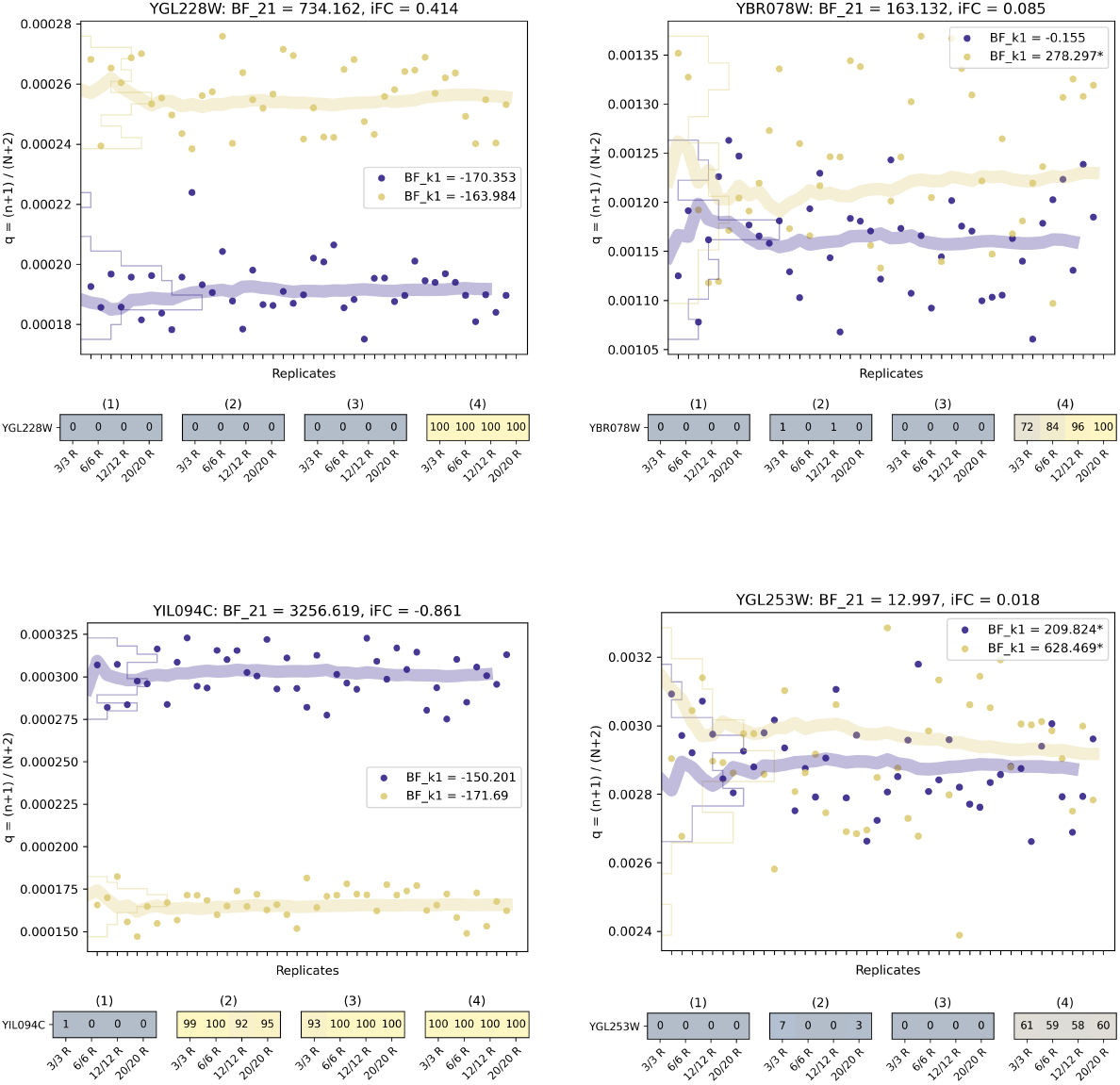
A set of example genes in yeast for which the DEG classification is sensitive to log_2_ fold change cut-offs. The inferred fold change in the Bayesian framework is consistently higher than the fold change calculated by *DESeq2* and *edgeR*, which causes classification differences for genes like YIL094C. The plots at the top show inferred expression probability, *q* = (*n* + 1)*/*(*N* + 2) with *n* reads mapping to the gene of interest and *N* total reads in the experiment. We show one gene per plot across 42 WT and 44 SNF2-mutant bulk, yeast culture RNA-Seq replicates [1]. The WT is shown in purple, and the SNF2-mutant in yellow. Note the different scales of the y-axes. The fine lines are density histograms obtained by projecting the points on the y-axis, and the thick lines are estimated means of *q*, updated with each replicate. For each gene Bayes factor (*BF*_21_) and inferred log_2_ fold change (*iFC*) over all replicates are given at the top. The bottom plot shows the number of times (out of 100) the gene has been identified as a DEG, for randomly sampled datasets with 3, 6, 12 and 20 replicates. We examine four DEG criteria: (1) *BF*_21_ > 1, and inferred | *iFC* | > 1; (2) *edgeR*: p-value *<* 0.05 and | log_2_ fold change | > 1; (3) *DESeq2* : p-value *<* 0.05 and | log_2_ fold change | > 1; (4) *BF*_21_ > 1, no log_2_ fold change cut-off. The (log) Bayes factors for expression consistency, *BF*_*k*1_, are given as insets in boxes [5]. If the gene is marked* it has been identified as ‘consistently inconsistent’, see Results section and Figure 3.

**Figure 6:**
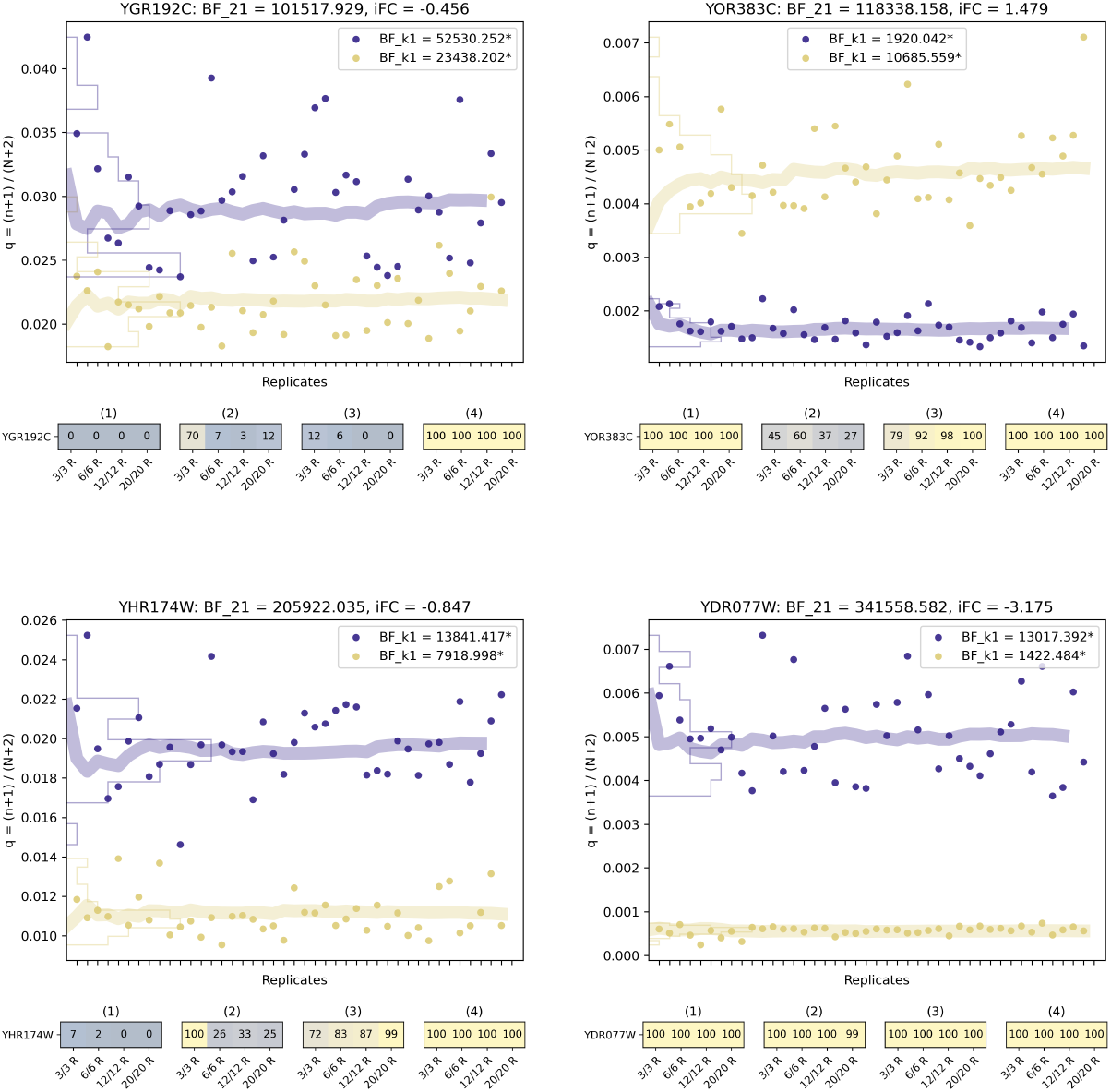
Example genes with some of the highest Bayes factors (*BF*_21_) over all replicates that are not always identified as DEGs. The plots at the top show inferred expression probability, *q* = (*n*+1)*/*(*N* +2) with *n* reads mapping to the gene of interest and *N* total reads in the experiment. We show one gene per plot across 42 WT and 44 SNF2-mutant bulk, yeast culture RNA-Seq replicates [1]. The WT is shown in purple, and the SNF2-mutant in yellow. Note the different scales of the y-axes. The fine lines are density histograms obtained by projecting the points on the y-axis, and the thick lines are estimated means of *q*, updated with each replicate. For each gene Bayes factor (*BF*_21_) and inferred log_2_ fold change (*iFC*) over all replicates are given at the top. The bottom plot shows the number of times (out of 100) the gene has been identified as a DEG, for randomly sampled datasets with 3, 6, 12 and 20 replicates. We examine four DEG criteria: (1) *BF*_21_ > 1, and inferred | *iFC* | > 1; (2) *edgeR*: p-value *<* 0.05 and | log_2_ fold change | > 1; (3) *DESeq2* : p-value *<* 0.05 and | log_2_ fold change | > 1; (4) *BF*_21_ > 1, no log_2_ fold change cut-off. The (log) Bayes factors for expression consistency, *BF*_*k*1_, are given as insets in boxes [5]. If the gene is marked* it has been identified as ‘consistently inconsistent’, see Results section and Figure 3.

**Figure 7:**
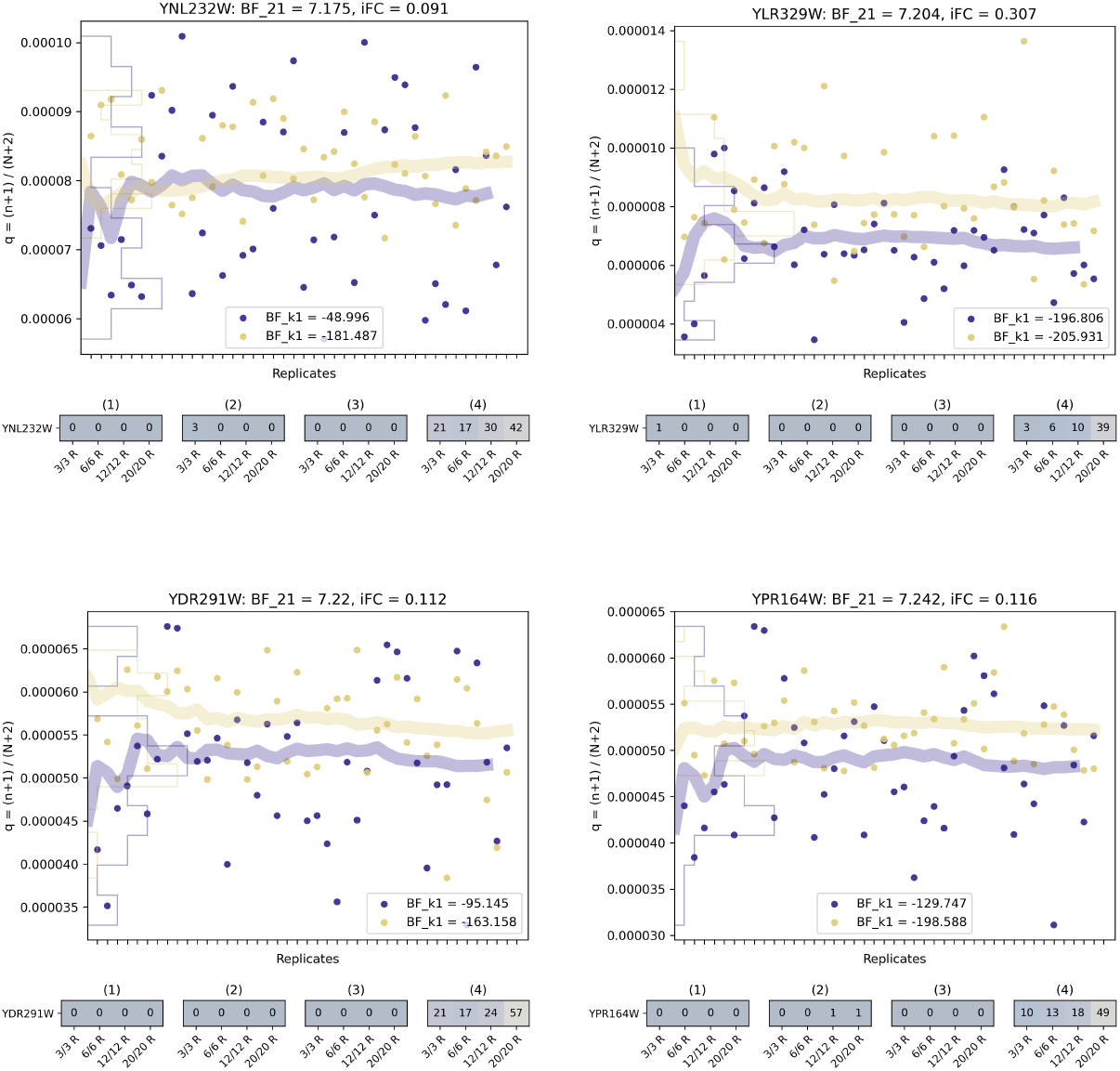
Different analyses do agree in the identification of no change. Small changes result in positive Bayes factors if a high number of replicates is given. The plots at the top show inferred expression probability, *q* = (*n* + 1)*/*(*N* + 2) with *n* reads mapping to the gene of interest and *N* total reads in the experiment. We show one gene per plot across 42 WT and 44 SNF2-mutant bulk, yeast culture RNA-Seq replicates [1]. The WT is shown in purple, and the SNF2-mutant in yellow. Note the different scales of the y-axes. The fine lines are density histograms obtained by projecting the points on the y-axis, and the thick lines are estimated means of *q*, updated with each replicate. For each gene Bayes factor (*BF*_21_) and inferred log_2_ fold change (*iFC*) over all replicates are given at the top. The bottom plot shows the number of times (out of 100) the gene has been identified as a DEG, for randomly sampled datasets with 3, 6, 12 and 20 replicates. We examine four DEG criteria: (1) *BF*_21_ > 1, and inferred | *iFC* | > 1; (2) *edgeR*: p-value *<* 0.05 and | log_2_ fold change | > 1; (3) *DESeq2* : p-value *<* 0.05 and | log_2_ fold change | > 1; (4) *BF*_21_ > 1, no log_2_ fold change cut-off. The (log) Bayes factors for expression consistency, *BF*_*k*1_, are given as insets in boxes [5]. If the gene is marked* it has been identified as ‘consistently inconsistent’, see Results section and Figure 3.

**Figure 8:**
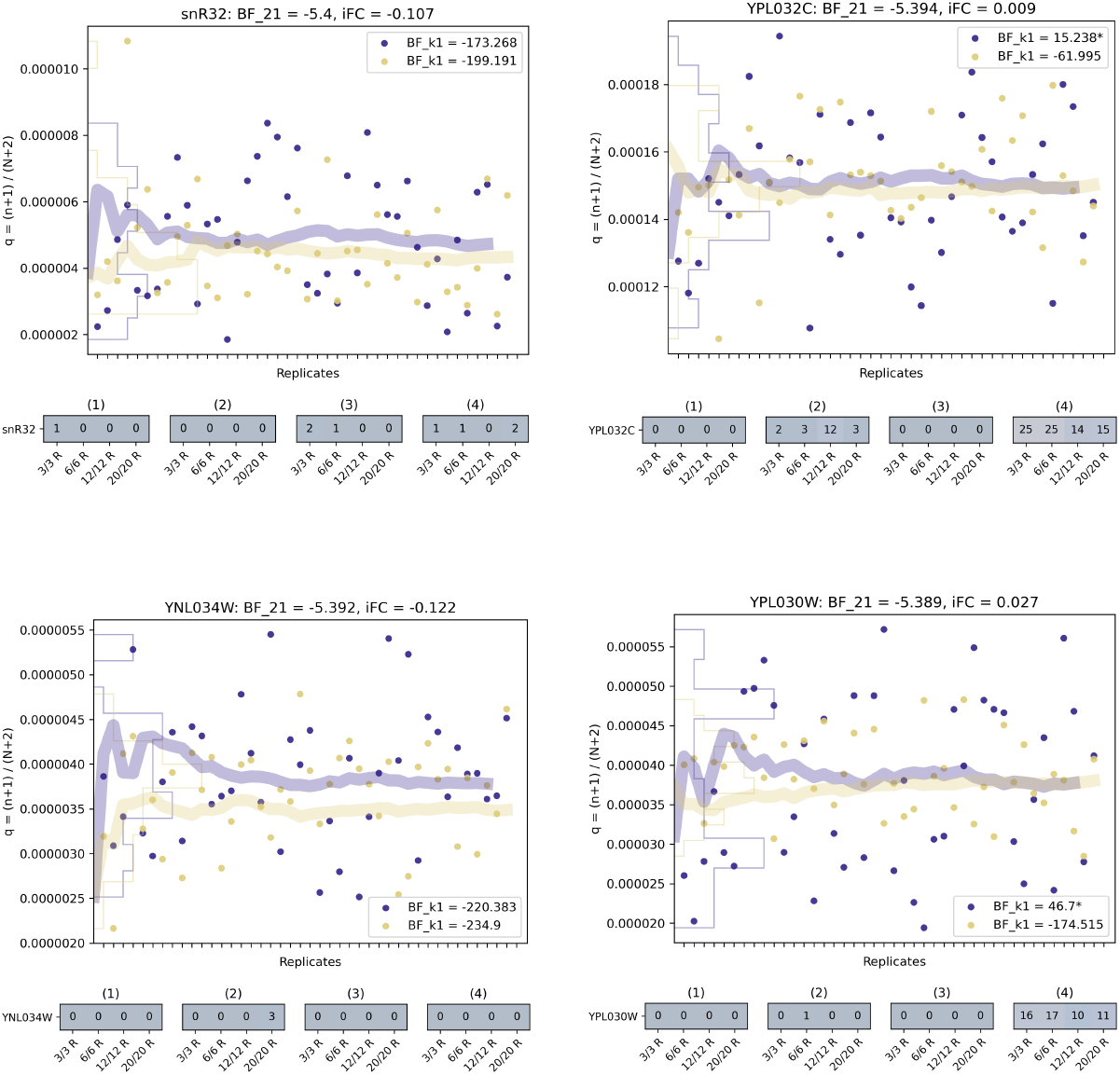
Four example genes with negative Bayes factors (*BF*_21_) over all replicates. The plots at the top show inferred expression probability, *q* = (*n* + 1)*/*(*N* + 2) with *n* reads mapping to the gene of interest and *N* total reads in the experiment. We show one gene per plot across 42 WT and 44 SNF2-mutant bulk, yeast culture RNA-Seq replicates [1]. The WT is shown in purple, and the SNF2-mutant in yellow. Note the different scales of the y-axes. The fine lines are density histograms obtained by projecting the points on the y-axis, and the thick lines are estimated means of *q*, updated with each replicate. For each gene Bayes factor (*BF*_21_) and inferred log_2_ fold change (*iFC*) over all replicates are given at the top. The bottom plot shows the number of times (out of 100) the gene has been identified as a DEG, for randomly sampled datasets with 3, 6, 12 and 20 replicates. We examine four DEG criteria: (1) *BF*_21_ >1, and inferred | *iFC* | > 1; (2) *edgeR*: p-value *<* 0.05 and | log_2_ fold change | > 1; (3) *DESeq2* : p-value *<* 0.05 and | log_2_ fold change | > 1; (4) *BF*_21_ > 1, no log_2_ fold change cut-off. The (log) Bayes factors for expression consistency, *BF*_*k*1_, are given as insets in boxes [5]. If the gene is marked* it has been identified as ‘consistently inconsistent’, see Results section and Figure 3.

**Figure 9:**
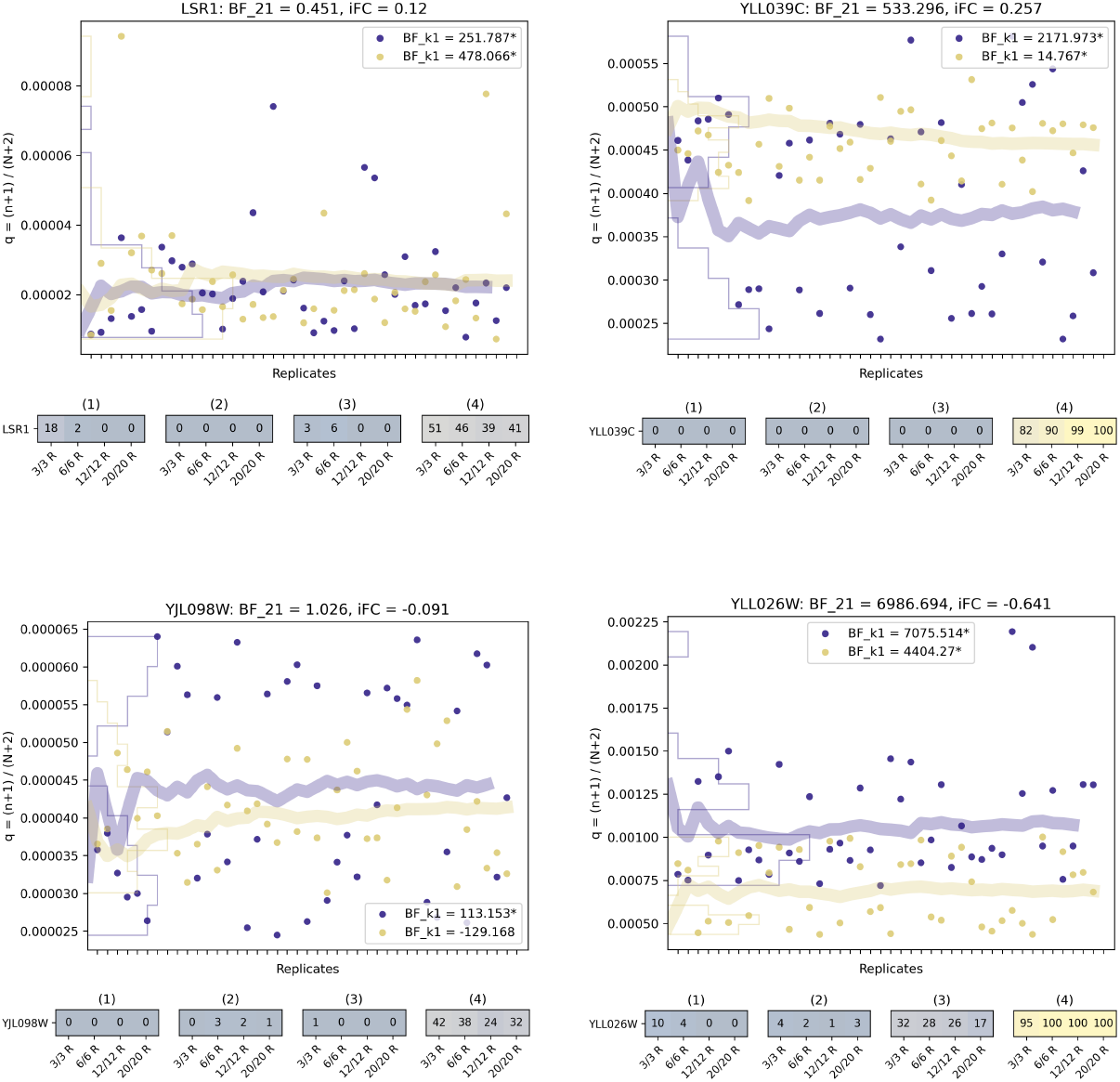
A set of consistently inconsistent genes that we identified in the Results section, Figure 3. The plots at the top show inferred expression probability, *q* = (*n* + 1)*/*(*N* + 2) with *n* reads mapping to the gene of interest and *N* total reads in the experiment. We show one gene per plot across 42 WT and 44 SNF2-mutant bulk, yeast culture RNA-Seq replicates [1]. The WT is shown in purple, and the SNF2-mutant in yellow. Note the different scales of the y-axes. The fine lines are density histograms obtained by projecting the points on the y-axis, and the thick lines are estimated means of *q*, updated with each replicate. For each gene Bayes factor (*BF*_21_) and inferred log_2_ fold change (*iFC*) over all replicates are given at the top. The bottom plot shows the number of times (out of 100) the gene has been identified as a DEG, for randomly sampled datasets with 3, 6, 12 and 20 replicates. We examine four DEG criteria: (1) *BF*_21_ > 1, and inferred | *iFC* | > 1; (2) *edgeR*: p-value *<* 0.05 and | log_2_ fold change | > 1; (3) *DESeq2* : p-value *<* 0.05 and | log_2_ fold change | > 1; (4) *BF*_21_ > 1, no log_2_ fold change cut-off. The (log) Bayes factors for expression consistency, *BF*_*k*1_, are given as insets in boxes [5]. If the gene is marked* it has been identified as ‘consistently inconsistent’, see Results section and Figure 3.

### 4.3 Identifying consistently inconsistent genes

Previously, we described how to calculate Bayes factors (*BF*_*k*1_) to test the consistency of replicates and rank genes according to their variability between replicates [5]. In order to identify highly variable genes in the yeast data set [1] we calculated *BF*_*k*1_ for all genes. To identify what we termed ‘consistently inconsistent’ genes, we used our randomly sampled datasets, and counted how many of the 7126 genes in yeast are identified as inconsistent, while increasing the number of replicates. For a gene to be considered what we term ‘consistently inconsistent’ we require *BF*_*k*1_ > 1, after analysing *k* replicates. The criterion had to be fulfilled in at least one of the 100 sampled datasets.

## Acknowledgements

We thank our colleagues Ander Movilla-Miangolarra and Pirita Paajanen for their comments and critical discussions on the manuscript. Furthermore, we would like to mention Emma Raven, Marina Millán-Blanquez, Burkhard Steuernagel, Philippa Borrill, Marek Glombik, Christine Faulkner and Hugh Woolfenden for many insightful conversations on the topic over the last years. Finally, we thank the JIC and TSL community – above all, the members of the Morris Lab – for thought-provoking discussions after seminars that shaped this work.

## 5 Appendix

### Differential Gene Expression Plot Collection

In previous work, we conducted a comparison between popular methods to identify differentially expressed genes [5], following similar studies by others [1]–[4]. We used popular packages to carry out differential expression analysis on a yeast data set and discussed the results [5]. Comparing the new Bayesian framework *bayexpress*, with *DESeq2* [4], [17] and *edgeR* [18], [26], we found discrepancies in the results. Here, after gaining new insights from our investigation into the effect of the number of replicates, we discuss the disagreements further. In the ‘Differential Gene Expression Plot Collection’ we illustrate instances where frameworks to identify DEGs do not agree.

We present a series of plots for single genes. At the top, we show the inferred expression probability, *q* = (*n* + 1)*/*(*N* + 2) with *n* reads mapping to the gene of interest and *N* total reads in the experiment, for 42 WT and 44 SNF2-mutant yeast RNA-Seq replicates. The plots below are the results of a comparison of three statistical frameworks: *bayexpress* [5], *DESeq2* [4], [17], and *edgeR* [18], [26]). We conducted an experiment to evaluate the impact of the number of replicates on the identification of DEGs. We generated 100 datasets by sampling 3, 6, 12, or 20 replicates from a pool of 42 WT and 44 SNF2-mutant replicates and performed differential gene expression analysis using the different methods on those 100 datasets. The numbers (and colours) are the counts out of 100 for how often the gene has been labelled as a DEG. We applied four different criteria: (1) BF_21_ > 1 and inferred | log_2_ fold change | > 1; (2) *edgeR*: p-value *<* 0.05 and | log_2_ fold change | > 1; (3) *DESeq2* : p-value *<* 0.05 and | log_2_ fold change | > 1; (4) *BF*_21_ > 1 and no inferred log_2_ fold change cut-off.

The first set of examples (Figure 4: ‘YAL016C-B’, ‘YAL031W-A’, ‘YAR035W’, ‘YAR068W’) was taken from a pool of genes that were identified as DEGs by Bayes factors (*BF*_21_), but not the other two packages. These genes are not highly expressed but show clear expression changes between WT and mutant. However, they are not reliably identified by *edgeR* (2) and *DESeq2* (3), even if 20 replicates are used. Bayes factors tend to increase with the number of replicates, meaning that DEGs are more often identified for higher numbers of replicates. The reasons for the classification differences between the frameworks are slight variations in fold change values between the packages, leading to them being just above or below the threshold. We compared the three fold change values and found that inferred log_2_ fold change values are consistently slightly lower, leading to classification differences [5].

The second set of examples (Figure 5: ‘YGL228W’, ‘YBR078W’, ‘YIL094C’, ‘YGL253W’) was chosen to determine the influence of fold change cut-offs. None of these genes were reliably classified as DEGs, despite noticeable changes. YIL094C is an example from a group of genes that are identified as DEGs by *DESeq2* and *edgeR* but not the Bayesian framework. The reason is the difference in how exactly log_2_ fold change values are calculated.

The third set of genes (Figure 6: ‘YGR192C’, ‘YOR383C’, ‘YHR174W’, ‘YDR077W’) shows some of the highest Bayes factors (*BF*_21_ across all replicates), presenting clear separations between WT and mutant expression. Note, that *BF*_21_ grows to astronomical numbers when summing up the evidence over 42/44 replicates, in line with increasing amounts of data supporting a consistent inference. Nevertheless, some of these genes are not reliably classified as DEG when fold change cut-offs are introduced.

The fourth set of examples (Figure 7: ‘YNL232W’, ‘YLR329W’, ‘YDR291W’, ‘YPR164W’) shows that analyses successfully identify genes where the change is not extreme enough. Averaging over 42/44 replicates (thick lines) we can see distinct tendencies of the genes in WT and mutant, leading to positive Bayes factors (*BF*_21_). For a lower number of replicates *BF*_21_ we do not find supporting evidence for expression change. Nevertheless, we can see again, how variability paired with low replication numbers can be misinterpreted as expression change.

The fifth set of examples (Figure 8: ‘snR32’, ‘YPL032C’, ‘YNL034W’, ‘YPL030W’) presents genes with negative *BF*_21_. Note, that in one of them the thick lines (estimated averages) are separated but we do not get a positive Bayes factor because there is not enough evidence given in the data, as there are not many reads mapping to this gene (see y-axis scale).

The last set of examples are consistently inconsistent genes that we identified in the Results section.

To summarise the findings of our framework comparison, we see some agreements but also interesting differences. For high number of replicates, the disagreements can be explained by arbitrary cut-offs, or filtering of genes with 0 reads [5]. For low numbers of replicates, the role of variability gains importance and may result in poor reproducibility. Both scenarios support our previous suggestion to renounce binary classifications and introduce rank-based methods [5].

### Supplementary Material

**Figure 10:**
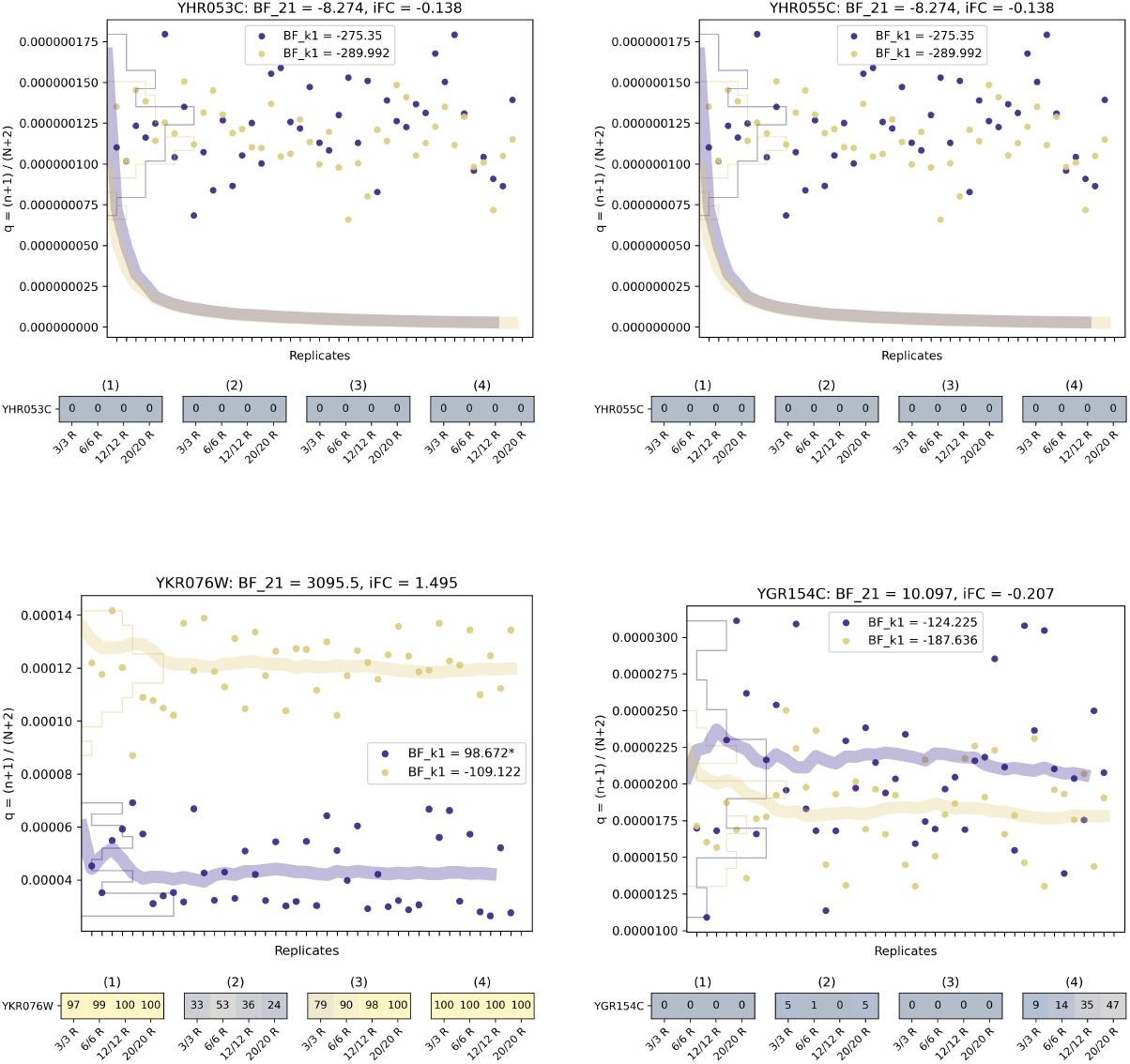
SNF2-targets (Table 1) and their gene expression change between wild-type and SNF2-mutant yeast (1/3). The bottom plot shows the number of times (out of 100) the gene was identified as a DEG, for randomly sampled datasets with 3, 6, 12 and 20 replicates. We examine four DEG criteria: (1) *BF*_21_ > 1, and inferred | *iFC* | > 1; (2) *edgeR*: p-value *<* 0.05 and | log_2_ fold change | > 1; (3) *DESeq2* : p-value *<* 0.05 and | log_2_ fold change | > 1; (4) *BF*_21_ > 1, no log_2_ fold change cut-off. The Bayes factors in boxes are *BF*_*k*1_ for testing the consistency of the number of mapping reads across replicates [5]. If the gene is marked* it has been identified as ‘consistently inconsistent’, see Results section and Figure 3. For a more detailed explanation and legend of the plots see Figure 1 or the Differential Gene Expression Plot Collection in the Appendix.

**Figure 11:**
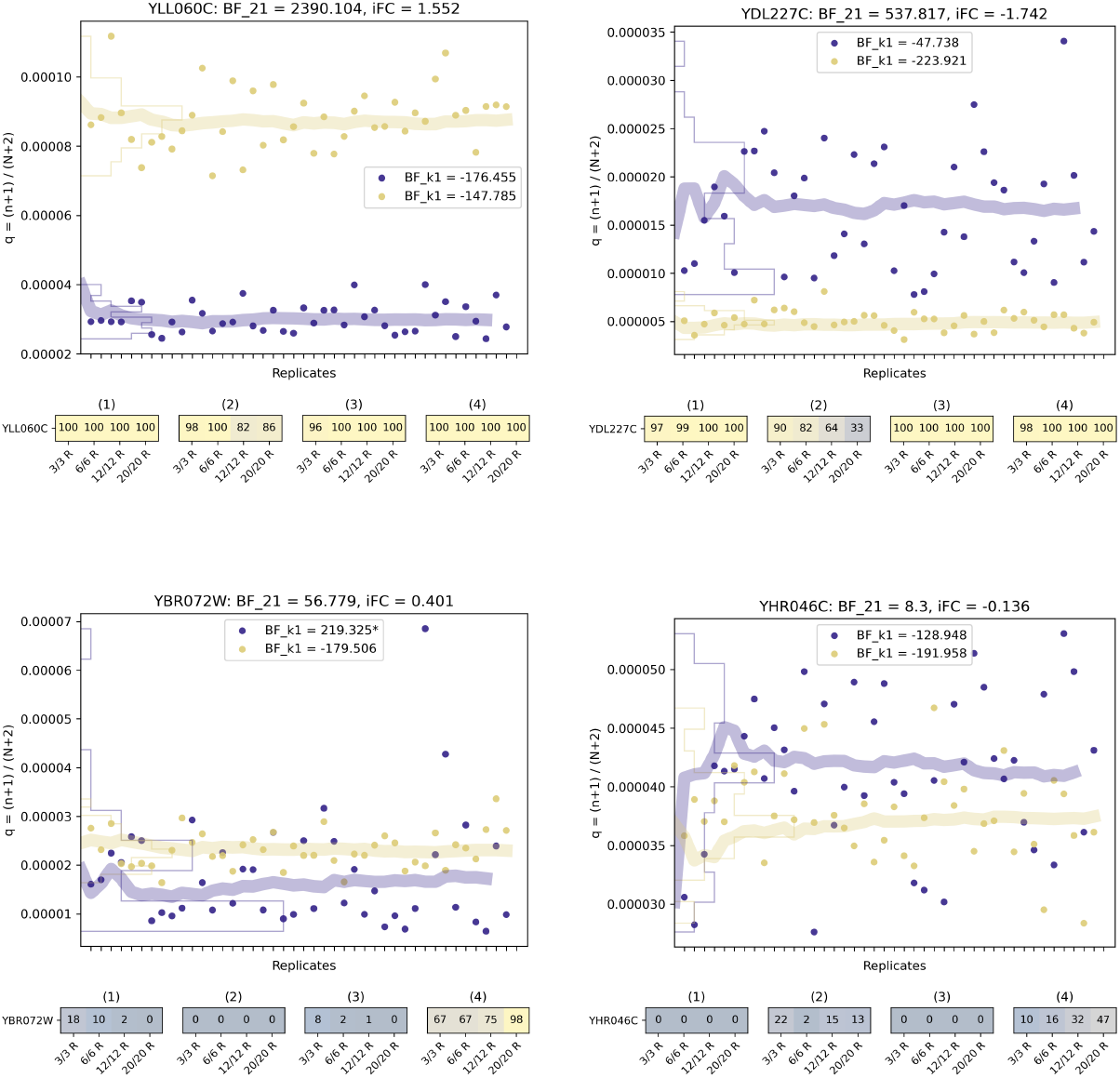
SNF2-targets (Table 1) and their gene expression change between wild-type and SNF2-mutant yeast (2/3). The bottom plot shows the number of times (out of 100) the gene was identified as a DEG, for randomly sampled datasets with 3, 6, 12 and 20 replicates. We examine four DEG criteria: (1) *BF*_21_ > 1, and inferred | *iFC* | > 1; (2) *edgeR*: p-value *<* 0.05 and | log_2_ fold change | > 1; (3) *DESeq2* : p-value *<* 0.05 and | log_2_ fold change | > 1; (4) *BF*_21_ > 1, no log_2_ fold change cut-off. The Bayes factors in boxes are *BF*_*k*1_ for testing the consistency of the number of mapping reads across replicates [5]. If the gene is marked* it has been identified as ‘consistently inconsistent’, see Results section and Figure 3. For a more detailed explanation and legend of the plots see Figure 1 or the Differential Gene Expression Plot Collection in the Appendix.

**Figure 12:**
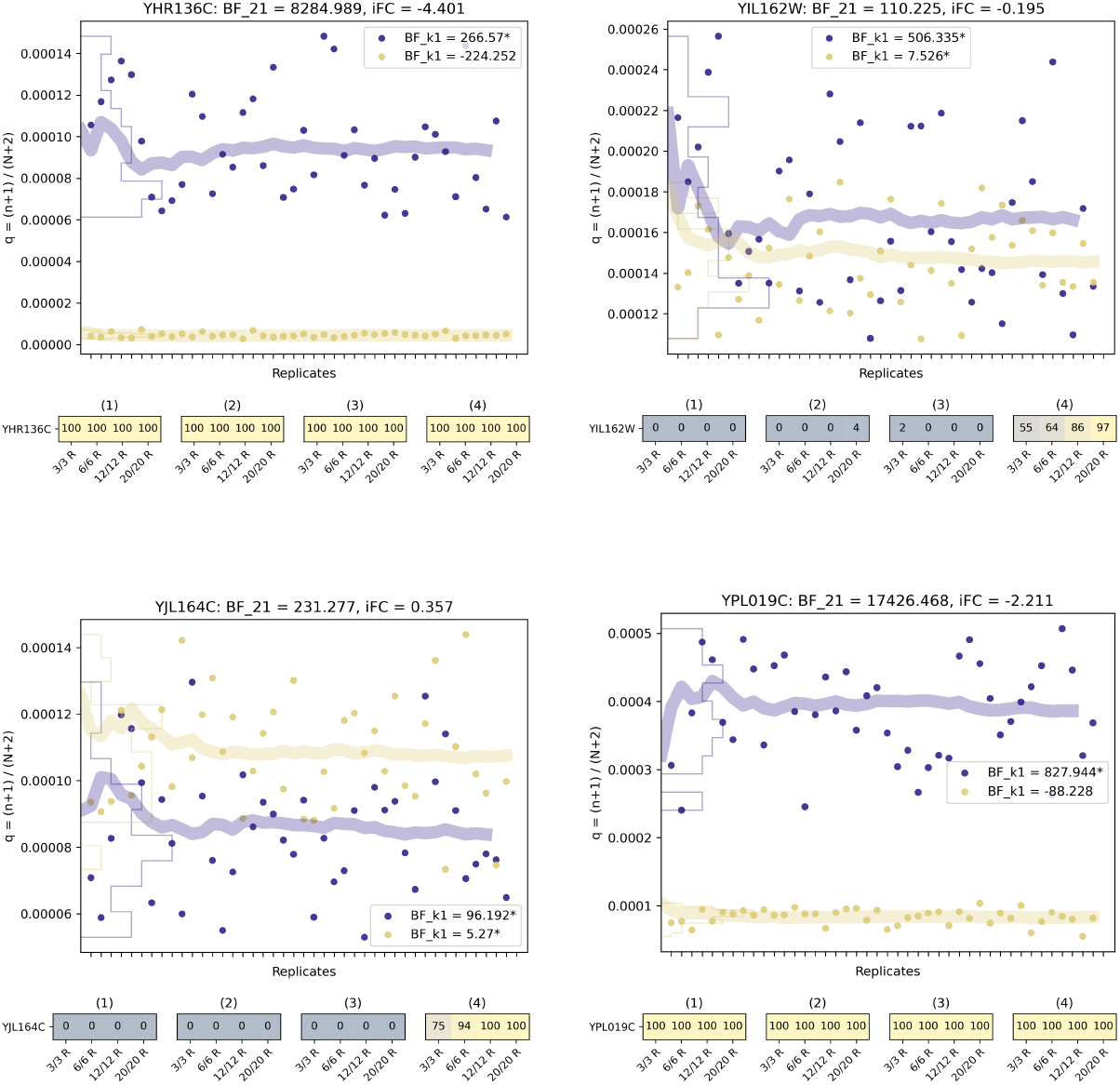
SNF2-targets (Table 1) and their gene expression change between wild-type and SNF2-mutant yeast (3/3). The bottom plot shows the number of times (out of 100) the gene was identified as a DEG, for randomly sampled datasets with 3, 6, 12 and 20 replicates. We examine four DEG criteria: (1) *BF*_21_ > 1, and inferred | *iFC* | > 1; (2) *edgeR*: p-value *<* 0.05 and | log_2_ fold change | > 1; (3) *DESeq2* : p-value *<* 0.05 and | log_2_ fold change | > 1; (4) *BF*_21_ > 1, no log_2_ fold change cut-off. The Bayes factors in boxes are *BF*_*k*1_ for testing the consistency of the number of mapping reads across replicates [5]. If the gene is marked* it has been identified as ‘consistently inconsistent’, see Results section and Figure 3. For a more detailed explanation and legend of the plots see Figure 1 or the Differential Gene Expression Plot Collection in the Appendix.

**Figure 13:**
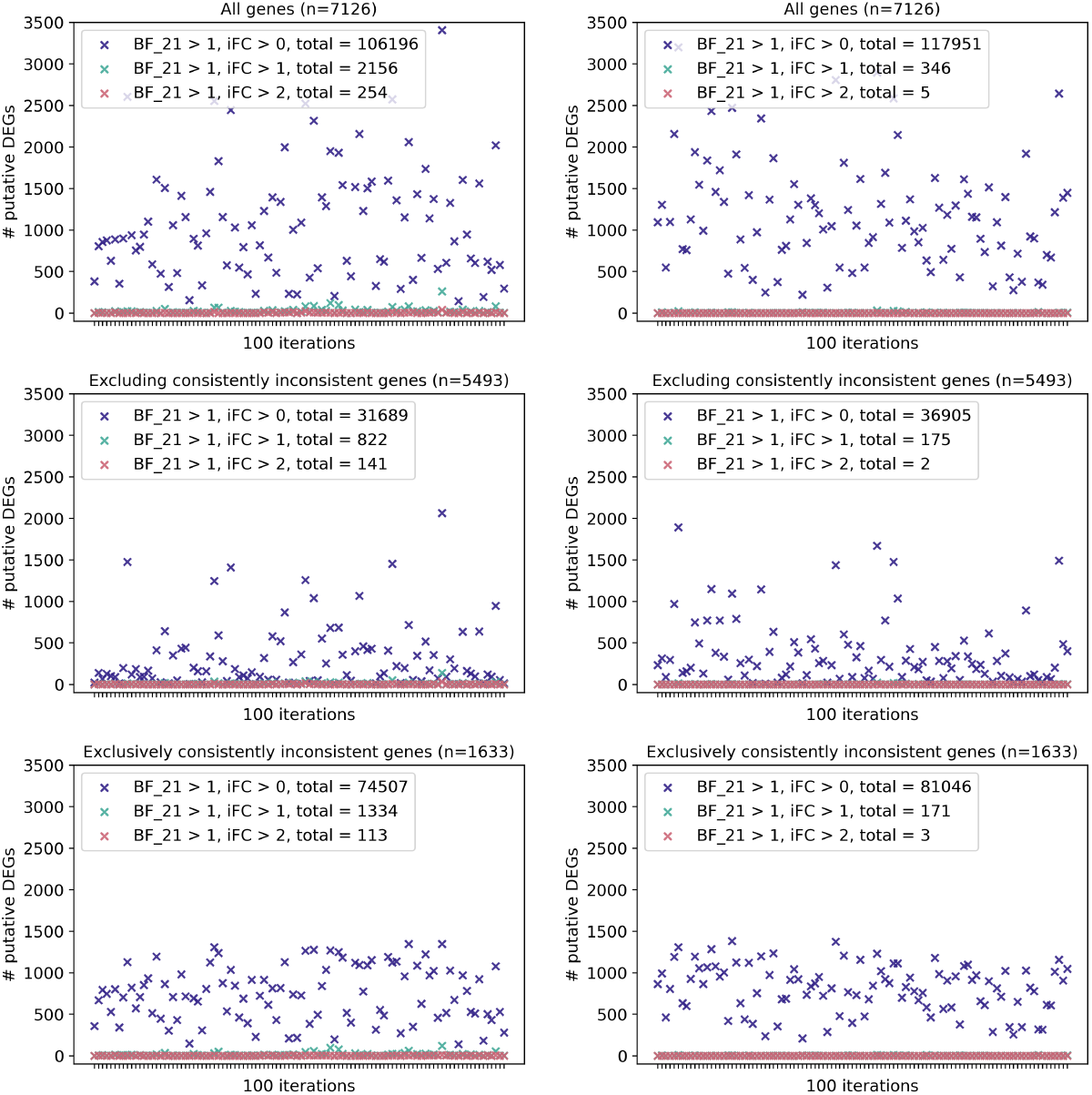
Control experiment (WT vs. WT): exploring the impact consistently inconsistent genes. Here, we show a control experiment comparing WT with WT yeast data to search for alleged differentially expressed genes with 3 different subsets of the genome. In the top plots, the whole genome (n=7126) is depicted, compare Results section. The middle plot depicts a set of genes that excludes genes that were identified as ‘consistently inconsistent’ (definition in Results and Methods). Considering three replicates (left column) and different log_2_ fold change cut-offs we lose 70% (purple: *BF >* 1, |*FC*| > 0), 62% (green: *BF >* 1, |*FC*| > 1) and 44% (red: *BF >* 1, |*FC*| > 2) of the false positives. Considering ten replicates (right column) and different criteria we are losing 69% (purple: *BF >* 1, |*FC*| > 0), 49% (green: *BF >* 1, |*FC* |> 1), and 3 out of 5 (red: *BF >* 1, |*FC*| > 2) of the false positives. The control experiment on the opposing set of genes, only considering the 1633 consistently inconsistent genes, is shown in the bottom plots.

